# Multimodal MEG and microstructure-MRI investigations of the human hippocampal scene network

**DOI:** 10.1101/2024.07.16.603546

**Authors:** Marie Lucie-Read, Carl J. Hodgetts, Andrew D. Lawrence, C. John Evans, Krish D. Singh, Katja Umla-Runge, Kim S. Graham

## Abstract

Although several studies have demonstrated that perceptual discrimination of complex scenes relies on an extended hippocampal network, distinct from an anterotemporal network supporting the perceptual discrimination of faces, we currently have limited insight into the specific functional and structural properties of these networks. Here, combining electrophysiological (magnetoencephalography, MEG) and microstructural (multi-shell diffusion MRI, dMRI) imaging in healthy human adults (30 female/10 male), we show that both hippocampal theta power modulation and fibre restriction of the fornix (a major input/output pathway of the hippocampus) independently related to accuracy during scene, but not face, perceptual discrimination. Conversely, microstructural features of the inferior longitudinal fasciculus (a long-range occipito- anterotemporal tract) correlated with face, but not scene, perceptual discrimination accuracy. Our results provide new mechanistic insight into the neurocognitive systems underpinning complex scene and face perception, providing support for multiple-system representation-based accounts of the medial temporal lobe.

**Significance Statement:** In contrast to theories positing segregated cortical areas for perception and memory, the specialized representations of the hippocampus may support both the perception and memory of visual scenes. To investigate, we utilised the unique window into hippocampal electrophysiological activity offered by Magnetoencephalography (MEG). We found hippocampal theta activity modulations in the hippocampus and posteromedial cortex during scene, versus face and shape-size, perceptual oddity discrimination, the magnitude of which correlated with scene, but not face or shape-size, discrimination accuracy. Moreover, multimodal white matter imaging revealed that tissue restriction of the fornix – the major hippocampal output tract - independently predicted scene discrimination performance. Our multimodal MEG-microstructure study provides novel evidence that the hippocampus and connected structures conjointly support online scene processing.

## Introduction

Challenging the long-standing view that the hippocampus exclusively supports long-term declarative memory, to the exclusion of other cognitive capacities (Squire & Dede, 2015), neuroimaging and lesion studies suggest that the hippocampus is better understood in terms of its role in forming complex scene (or relational) representations that support episodic memory, but also ‘on-line’ processing including complex scene discrimination (Gardette et al., 2022; Graham et al., 2010; Hodgetts, Voets, et al., 2017; Lee, Buckley, et al., 2005; Murray et al., 2017; Ruiz et al., 2020; Zeidman & Maguire, 2016). Critically, the hippocampus’s role in scene representation is thought to emerge primarily through interactions within an “extended hippocampal” or “posteromedial” system - comprising the hippocampus, mammillary bodies, anterior thalamic nuclei and retrosplenial cortex (Aggleton, 2012; Gaffan & Hornak, 1997; Murray et al., 2017; Ranganath & Ritchey, 2012). Indeed, by imaging inter-individual differences in brain microstructure with diffusion magnetic resonance imaging (dMRI) alongside performance of an ‘odd-one-out’ or oddity perceptual discrimination task, we previously found that inter-individual variation in microstructure (mean diffusivity, MD) of the fornix – a tract that interconnects regions within the extended hippocampal system (Aggleton, 2012) – was related to the ability to discriminate scenes, but not faces (Hodgetts et al., 2015). By contrast, inter-individual microstructure variation of the inferior longitudinal fasciculus (ILF) (Herbet et al., 2018), a ventral long-range fibre tract interconnecting the occipital and anterior temporal lobes, correlated with face, but not scene, discrimination. We also found evidence for associations between fornix and ILF microstructure and category-selective blood oxygen-level dependent (BOLD) responses in the hippocampus and the fusiform face area and perirhinal cortex for scenes and faces, respectively.

Despite these findings, given the indirect link between the BOLD signal and neural activity (Ekstrom, 2021), we lack understanding of specific neurobiological mechanisms that support information processing within this extended hippocampal system, and therefore scene discrimination. Magnetoencephalography (MEG) offers a unique window into the behavioural correlates of hippocampal electrophysiological activity (Alberto et al., 2021; Ruzich et al., 2019). Notably, Barry et al. (2019) have recently shown modulation of hippocampal theta (∼4-8 Hz) oscillatory activity during the construction of novel scene imagery. Specifically, scene imagination was associated with *attenuation* of theta oscillatory power (see also Monk et al., 2021).

To investigate whether such findings extend to a role for the hippocampus in perception, we recorded theta power modulations, using MEG, in participants performing a perceptual ‘odd-one-out’ discrimination task for scenes and faces (plus a difficult size oddity control condition). We predicted that theta power modulations for scenes (relative to non-scene conditions) would occur across the hippocampus and connected regions within the extended hippocampal system, including posteromedial regions encompassing retrosplenial cortex (Aggleton, 2012; Baldassano et al., 2016; Hodgetts et al., 2016). We then examined whether individual differences in hippocampal theta modulation would be related to scene, but not face, oddity discrimination abilities.

We next investigated the relationship between oddity performance and specific tissue properties of the fornix and ILF, alongside the parahippocampal cingulum bundle (PHCB) (Bubb et al., 2018). Tissue properties were obtained from principal components analysis (PCA)-based reduction of multiple indices derived from advanced biophysical (Neurite Orientation and Dispersion Density Imaging, NODDI; Composite Hindered And Restricted Model of Diffusion, CHARMED) and standard tensor-derived (DTI) models applied to multi-shell dMRI data, together with indices derived from quantitative Magnetization Transfer (qMT) imaging (Chamberland et al., 2019; Read et al., 2023). We predicted that tissue properties of the fornix would be associated with scene, but not face oddity discrimination, and vice versa for the ILF.

Finally, based on the proposed link between regional functional specialization and long-range structural connectivity (Chen et al., 2017) and evidence of a link between (developmental) variations in white-matter tissue properties, electrophysiological responses, and behaviour (Caffarra et al., 2024), we examined potential three-way relationships between hippocampal theta power modulation, white-matter microstructure, and scene oddity discrimination. Our results provide new insights into the role of an extended hippocampal system in complex scene perception.

## Materials and methods

### Participants

Forty-three adult volunteers, with no reported neurological pathology, participated in a MEG session with cognitive tasks, and a Magnetic Resonance Imaging (MRI) session within a fortnight of each other. Due to data collection disruptions, data from three participants were incomplete, leaving forty MEG-oddity datasets (mean age: 22.5 years, SD 4.0, range: 18-38 years; 30 female/10 male). Of these, one participant requested to leave the MRI scans early, resulting in thirty-nine microstructure datasets (mean age: 22.5 years, SD 4.2, range: 18-38 years; 29 female/10 male). This study was approved by the Cardiff University School of Psychology Research Ethics Committee and all participants provided written informed consent.

### Oddity task and procedure

The cognitive tasks were presented using Psychtoolbox (Kleiner et al., 2003) for MATLAB (MATLAB, 2015a). The oddity task was modified from previous fMRI studies (Barense et al., 2010; Hodgetts et al., 2015; Lee et al., 2013) for use in MEG. In this, participants examined simultaneously presented triplet images and identified the odd-one-out (Figure 1A). For the scene and face stimuli, the images were shown at three different angles and one image had either differing spatial object relationships (scene condition) or differing facial features (face condition). The control task (size condition) was designed to not require MTL-supported representations (Barense et al., 2010; Hodgetts et al., 2015). It comprised 3 circles shown with different locations within the three image spaces, with one circle differing in size (see Fig. 1A). In all conditions, the location of the odd image was counterbalanced across trials. Before scanning, participants practised the task with 8 trials of each condition. All stimuli were trial-unique.

**Figure 1.**
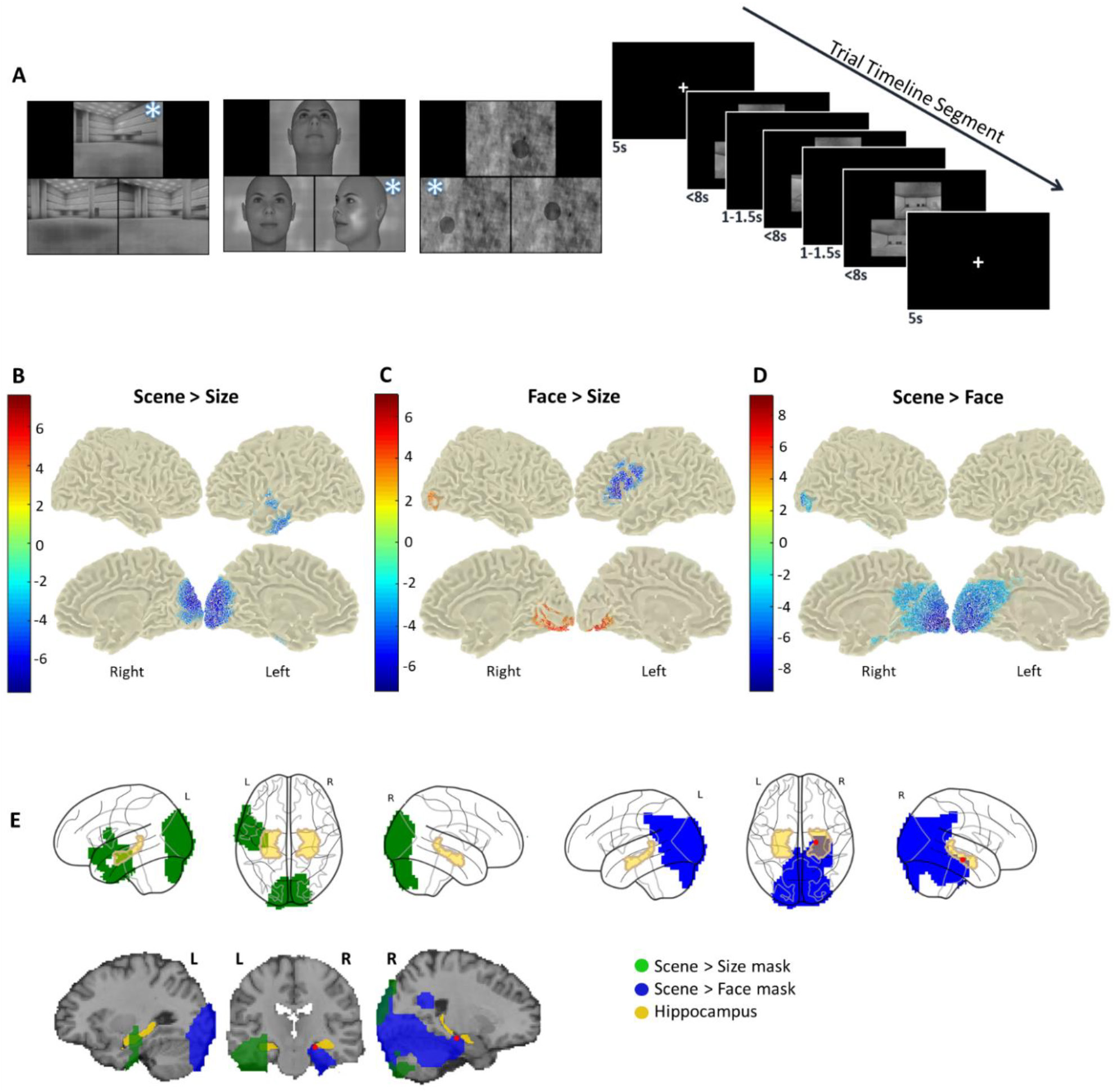
Oddity task outline and whole-brain theta power differences between conditions. **A**) Left: three trial examples. The triplets of images were presented simultaneously. For these examples, an asterisk is placed over the odd-one-out. Right: An illustration of a mini-block comprising three trials of the same condition, separated by short (1 - 1.5 s) inter-trial fixation periods, preceded and followed by 5 s fixation periods. The trials could be displayed up to 8 s in total but ended as soon as the participant made a response. **B**) scene - size comparison. **C**) face - size comparison. **D**) scene - face comparison. In each section, colours represent t-values according to the keys to the left, and the significant clusters are overlaid on template brain surfaces. **E**) The same masks of significant clusters for the scene>size and scene>face comparisons are overlaid (in green and blue, respectively) onto glass brains (separately) and a template T1 (together). For illustration, the hippocampus is shown in yellow (the Cornu Ammonis areas and subiculum probabilistic ROIs from the FSL Jülich histological atlas thresholded at 50%). The red dots indicate the location of the most negative t-value in the scene>face comparison (MNI coordinates: 22 −20 −16). Note that the alpha value was 0.017 (0.05 / 3 comparisons). L = Left. R = Right. Brain images made with Fieldtrip and Nilearn.

Participants responded using a button-box with their right hand and three keys represented the three images. Trials were sequentially displayed in mini-blocks of three trials of the same condition (Figure 1A). Mini-blocks were presented in a pseudo-random order, such that each mini-block’s condition was different from the previous one. The first trial of each mini-block was preceded by a 5 s fixation period of a black screen with a white cross at the centre (fixation condition). Other trials of the mini-block were preceded by an inter-trial fixation period lasting randomly between 1 s and 1.5 s. Participants were shown trials for up to 8 s but each trial ended as soon as a response was made, to reduce inclusion of post-decision mnemonic processes.

There was a total of 96 size trials and 144 face and scene trials each (plus an additional 8 practice trials of each). For each participant, the oddity task included all 96 size trials, 96 of the face trials, and 96 of the scene trials. To reduce fatigue and head movement over trials, the resulting 288 trials of the oddity task were split into four stimuli-counterbalanced blocks of 72 trials.

To test if hippocampal involvement during the oddity task was solely a reflection of incidental encoding, there was a subsequent surprise memory task, still during the MEG scan. Participants were presented with a series of 48 new scene and 48 new face trials plus 48 of the previously used scene trials and 48 of the previously used face trials (192 trials in total). Stimuli appearing in the oddity and memory tasks were counterbalanced across participants.

Participants responded using a button with their right hand and four keys represented four answers: ‘definitely old’, ‘I think it’s old’, ‘I think it’s new’ and ‘definitely new’. For this study, answers were grouped together and analysed as answered ‘old’ or ‘new’. Since decision making in the memory task was predicted to be faster than that of the oddity task, trials lasted for up to 3.5 s and mini-blocks were separated by a 2 s fixation period. As in the oddity task, the inter-trial fixation period randomly varied between 1 s and 1.5 s.

### Stimulus design

Most of the scene stimuli were generated by Lee et al. (2013), using Deus Ex (Ion Storm, 2000) and an editor (Deus Ex SDK), and were edited for this study (16 additional scenes were made using the same methods).

Face stimuli were generated using Facegen (Singular Inversions, 1998). After applying the default settings for face type (race, age, symmetry), setting the sex rating equidistant between male and female, and removing the facial hair, the ’generate random face’ function was used. The odd-one-out was constructed by using the ’genetics’ tab and applying a variation of 0.4. Faces within a trial were presented from three different viewpoints from four possibilities, ’right’ (45° right), ’left-up’ (45° left and 20° up), ’up’ (20° up) and straight-on (0°).

To minimise differences in image statistics between the task and control conditions, the size images were constructed using phase-scrambled versions of the scene and face images. Phase-scrambling was performed using a technique (Perry, 2016) that allows the user to determine the level of phase scrambling by using the ‘weighted mean phase’ method (Dakin et al., 2002). A weighting factor (determining the proportion of unaltered spatial phase in the scrambled image) of 0.16 was used because this has been shown to produce a subthreshold detection rate (Perry, 2016). Then, using the python package ‘PIL’ (Umesh, 2012), translucent homogenous black circles were placed over the scrambled images. Circles were randomly sized between 60x60 pixels to 90x90 pixels and one differed in size (pixel change of +/-4).

As visual signals can be stronger than those from deep brain sources, contrasting conditions with similar visual properties can be beneficial when measuring MTL signals with MEG (Quraan et al., 2011). To reduce differences in image statistics between conditions, all condition images were turned greyscale and altered using the SHINE toolbox for MATLAB (Willenbockel et al., 2010). Luminance was normalized across images using the ‘lumMatch’ function and the luminance histograms were matched using the ‘histMatch’ function. The Fourier amplitudes were matched across stimuli using ‘sfMatch’. To improve the quality of the histogram-matched images, ‘structural similarity index’ (Wang et al., 2004) was optimised over 15 iterations (Avanaki, 2009; Willenbockel et al., 2010).

### MEG recording and analyses

The MEG recordings were performed using a 275-channel (excluding 1 faulty channel) axial gradiometer CTF system, located inside a magnetically shielded room. The data were acquired continuously, with a sampling rate of 1200 Hz. Electromagnetic coils were placed on three fiducial locations, the nasion, and left and right pre-auricular regions. During the MEG recording, these sensors were energised with a high-frequency signal, to locate their positions relative to the MEG sensors. The locations of the fiducial points for each participant, and head shape, were recorded digitally using an Xsensor camera system (ANT Neuro, Enschede, The Netherlands), for subsequent co-registering to each participant’s T1-weighted MRI scans. To reduce the effects of head motion over long recordings, head localization was performed before each of the four recording blocks.

MEG analyses were carried out using the Fieldtrip toolbox (2019; Oostenveld et al., 2011) for MATLAB, with a pipeline based upon that of Magazzini & Singh (2018). First, the recordings were inspected manually for muscle and system artefacts before being downsampled (600Hz) and decomposed into 100 components using independent component analysis (with Fieldtrip’s fast ICA). Components relating to eye-movement, heart rate, and movement, were removed from the original data. These data were then cut into trials and visually inspected. Trials with remaining artefacts were manually excluded. In addition, an error meant that some participants viewed a scene image twice, so these trials were also removed.

We studied an early 2 s time period of the trial, starting at 0.3 s post stimulus onset, to exclude the majority of the visual evoked potentials which make deep brain source localization challenging (Quraan et al., 2011; see Rudoler et al., 2023 for a similar approach), and lasting until 2.3 s. As trials ended when participants responded, trial lengths varied, resulting in different average trial length between the conditions (see RTs in Results). Trials shorter than 2 s were not included in the analyses, which resulted in unequal trial numbers across conditions. After data cleaning and cutting, there were 82 scene trials, 82 face trials, 65 size trials and 88 fixation trials, per participant, on average.

#### Whole brain power

Data were downsampled to 300Hz, and condition trials were cut from 0.3-2.3 s and the fixation trials were cut from 1-3 s (to exclude post-stimulus effects). Source-localised modulations in the theta frequency band (4-8Hz) were calculated by band-pass filtering the data within this frequency band and carrying out source localisation, in the 2 s window of each trial, using the Linearly Constrained Minimum Variance beamforming method (LCMV: Van Veen et al., 1997).

LCMV beamforming was carried out by first estimating the covariance matrix across all trials to obtain common grid filters, and then applying these pre-computed spatial filters to each of the condition trials separately. To reduce the magnitude of participant movement in each beamformer calculation, oddity MEG recording blocks were analysed separately. The source images were then interpolated to an MNI template-space model (included with Fieldtrip), appended across MEG recording blocks, and anatomically labelled using the Automated Anatomical Labelling (AAL) atlas (Tzourio-Mazoyer et al., 2002). First, the conditions were compared at the individual level so that unequal trial numbers between the conditions could be accounted for. Average power within the 0.3-2.3 s trials were compared across conditions with Fieldtrip’s ‘ft_sourcestatistics’ using MATLAB’s t-test with unequal variance. This resulted in t-maps for each condition comparison (scene vs size, face vs size, scene vs face) for each participant. To implement a one-sample t-test across participants, of these within-participant effects, these condition effect t-maps were compared to equivalently-sized maps of zeros at the group level, using a paired-samples t-test with Monte Carlo sampling and 5000 permutations.

Using the resulting scene>face significant clusters mask and t-value map, minimum t-value (peak reductions) locations were identified in the bilateral hippocampus ROIs.

#### Hippocampal theta modulation from baseline

To measure individual differences in theta modulation from baseline, scene and face conditions were compared to the fixation condition. The same frequency and source analysis steps described above were carried out without statistical comparisons, but the average power modulations (across trials) were calculated as percentage changes between scene/face conditions and the fixation condition. The hippocampal mask was made using bilateral hippocampal AAL ROIs. Percentage power change within the voxels of this mask was averaged to create the hippocampal oscillatory power modulation values for each participant.

### MRI Recording and analyses

#### Scanning protocol

Structural MRI data were collected using a Siemens Prisma 3T MRI system with a 32- channel head coil. T1-weighted anatomical images were obtained using an MPRAGE sequence with the following parameters: slices = 176, time to repetition (TR) = 2300 ms, FOV = 256 mm x 256 mm, matrix size = 256 mm x 256 mm, flip angle = 9°, echo time (TE) = 3.06 ms, slice thickness = 1 mm.

Diffusion weighted data were acquired using a multi-shell HARDI protocol (Assaf & Basser, 2005; Santis et al., 2014) with the following parameters: phase encoding = A>P; slice thickness = 2 mm; TE = 73 ms; TR = 4100 ms; 203 gradient directions and 4 shells (maximum b-value: 4000 s/mm^2^); FOV = 220 mm x 220 mm. In addition, a reference acquisition with the opposite phase encoding direction (P>A) was acquired for blip-up blip-down correction, with 33 directions and 2 shells (maximum b-value: 1200 s/mm^2^) (Andersson et al., 2003; Andersson & Sotiropoulos, 2016).

MT-weighted data were acquired through an optimized 3D MT-weighted fast spoiled-gradient recalled-echo (SPGR) sequence (Cercignani & Alexander, 2006) with the following parameters: TR = 32 ms, TE = 2.46 ms, flip angle = 5°, bandwidth = 330 Hz/Px, FOV = 240 mm x 240 mm, slice thickness = 2 mm. The 11 MT-weighted volumes used Gaussian MT pulses of duration 12.8 ms the following with off-resonance irradiation frequencies/saturation pulse amplitudes: 1000 Hz/332°, 1000 Hz/333°, 12060 Hz/628°, 47180 Hz/628°, 56360 Hz/332°, 2750 Hz/628°, 1000 Hz/628°, 1000 Hz/628°, 2768 Hz/628°, 2790 Hz/628°, 2890 Hz/628°. In addition, maps of the RF transmit field (B_1_ ^+^) were collected using a pre-saturation TurboFLASH (satTFL) acquisition with the following parameters: TR = 5000 ms, TE = 1.83 ms, flip angle = 8°, matrix = 64 × 64. B0 maps were calculated using two gradient recalled acquisitions with the following parameters: TE = 4.92 ms and 7.38 ms; TR = 330 ms; FOV = 240 mm; slice thickness = 2.5 mm.

#### Diffusion MRI data processing

Diffusion data were denoised (Veraart et al., 2016). Motion distortion correction was carried out using the Eddy tool in FSL (Jenkinson et al., 2012). Full Fourier Gibbs ringing correction was carried out using mrdegibbs MRtrix software (Tournier et al., 2012). The separate contribution of the free water compartment to the DTI data was identified and removed from these data by a customized version of the Free Water Elimination (FWE-DTI) algorithm (Pasternak et al., 2009).

Tractography analysis was applied to the 1400 b-value shell. To detect and eliminate signal artefacts, the Robust Estimation in Spherical Deconvolution by Outlier Rejection (RESDORE) algorithm was applied (Parker et al., 2012). Subsequently, peaks in the fibre Orientation Distribution Function (fODF) in each voxel were extracted using the damped Richardson-Lucy technique (Dell’acqua et al., 2010). Whole-brain deterministic tractography was carried out in Explore DTI (version 4.8.3; Leemans et al., 2009). The streamlines were constructed using an fODF amplitude threshold of 0.05, step size of 0.5 mm and an angle threshold of 45°.

Tensor fitting was carried out on the 1200 b-value shell. To estimate the diffusion tensor in the presence of physiological noise and system-related artefacts, the Robust Diffusion Tensor Estimation (RESTORE) algorithm was applied (Chang et al., 2005). This analysis resulted in FA, MD and RD maps.

NODDI maps were created using the Accelerated Microstructure Imaging via Convex Optimization NODDI algorithm (Daducci et al., 2015). CHARMED analysis was carried out using an in-house program coded in MATLAB that calculated FR per voxel.

#### Magnetization transfer-weighted data

The magnetization transfer-weighted SPGR images were co-registered (affine, 12 degrees of freedom), within each participant, to the image with the highest contrast, to correct for interscan motion, using Elastix (Klein et al., 2010). Modelling was then carried out by using two-pool pulsed-magnetization transfer approximation as described by Ramani et al. (2002), which corrects for B and B ^+^ field inhomogeneities and produces MPF maps.

#### Tractography and tract microstructure features

Fornix, ILF and PHCB streamlines were generated using the protocols previously described in Read et al. (2023). In short, ’way-point’ ROIs were manually drawn onto whole-brain FA maps in the diffusion space of 18 subjects, using Explore DTI, to isolate individual tracts.

These streamlines were then used to train in-house automated tractography software (Greg Parker, Cardiff University; written in MATLAB, 2015), and the resulting tract models were then applied to the entire dataset.

FA, MD, RD, FR, MPF, NDI and OD values for the voxels encompassed in the tract streamlines were extracted and the mean was calculated for each tract. This resulted in seven microstructure metrics for three tracts for 39 participant datasets.

Microstructure data were reduced through PCA using the same methods as those described in Read et al. (2023). This approach has been shown to be effective in capturing biologically informative features in previous microstructure datasets (Chamberland et al., 2019; Geeraert et al., 2020; Henriques et al., 2023). In short, the Bartlett test was used to assess the data’s appropriateness for PCA, the prcmp function in R (R Core Team, 2019) was then used to apply PCA to centred and scaled data, and the sampling adequacy of the results was tested using the Kaiser-Meyer-Olkin (KMO) test (from the R ‘Psych’ package; Revelle, 2020).

Components were retained depending on the amount of cumulative variation they explained. Following data reduction, participant scores in two principal components (PCs) were used for analysis.

## Statistical analysis

Statistical analyses were carried out using Fieldtrip for MATLAB or using Rstudio and additional R packages (Kassambara, 2019; Oostenveld et al., 2011; Patil, 2021; R Core Team, 2019; R Studio Team, 2015; Wickham, 2016). Analysis Of Variance (ANOVA) tests and 95% confidence intervals were calculated in JASP (Version 0.18.1; JASP Team, 2023). Note that the 95% CIs reported throughout were derived using a 1000 iteration bootstrapping procedure.

The alpha thresholds for the whole-brain MEG condition power/connectivity comparisons were Bonferroni-corrected to 0.017 (0.05 / 3 comparisons). The cluster alpha threshold was 0.001. Fieldtrip’s ‘correct tail = alpha’ option was applied to further correct for two-sided tests.

Pearson’s correlation tests were applied to understand relationships between tract microstructure, oscillatory power and oddity performance. In cases where variable data did not have a normal distribution, the data were transformed (squared) to de-skew the distribution (McDonald, 2014). Coefficients of correlations were compared using the Pearson and Filson’s test within the R package ‘Cocor’ (Diedenhofen & Musch, 2015). Bayesian correlation tests were also calculated (Morey & Rouder, 2018) and Bayes Factors (BFs) were reported as BF10 (evidence of the alternative model over the null model).

Since Hodgetts et al. (2015) found negative correlations between fornix MD and scene oddity accuracy, and ILF MD and face oddity accuracy, and positive correlations between fornix FA and scene oddity accuracy, and ILF FA and face oddity accuracy, there were directed hypotheses about correlations between microstructure and oddity task performance. Therefore, the contribution of FA and MD values to microstructure PCA components, prescribed the hypotheses of how the components of the tracts would relate to oddity task performance, supporting the use of one-tailed statistical tests. Similarly, as the scene oddity task induced hippocampal theta attenuation, we predicted improved oddity performance with decreased hippocampal theta power, also supporting the use of one-tailed statistical tests.

Partial correlations were used for correlation analyses involving MEG data so that MEG trial numbers in each condition could be controlled for. MEG trials in which the participant did not respond were not included in the MEG analysis (to reduce the influence of off-task thoughts) but were regarded as incorrect in the behavioural data (as removing such trials would inflate performance). Although this is a small proportion of excluded MEG trials, there was still an association between the number of MEG trials and oddity performance. Therefore, partialling - out the variance from MEG trial numbers was required to adjust for its potential biasing of performance-MEG data correlations.

For the microstructure-oddity performance correlation tests, the alpha level was Bonferroni-corrected by dividing by the number of statistical comparisons involving each microstructure PC (0.05/3 oddity accuracy measures = 0.017; Hodgetts et al., 2015). This rule was also used for the oscillatory power-oddity performance correlation tests and oscillatory power-microstructure correlation tests since the alpha level was Bonferroni-corrected by dividing by the number of statistical comparisons involving each microstructure PC or oscillatory modulation measure. Similarly, when comparing correlation coefficients, the alpha level was Bonferroni-corrected by dividing by the number of statistical comparisons relating to a variable (0.05/2 = 0.025).

## Results

### Behavioural Results

Task difficulty (i.e., proportion correct) across the conditions was very well matched, as all condition accuracy means were around 61% (see Table 1). To reduce the scene accuracy data skew (to <1), so that parametric tests could be used while keeping the conditions matched, all accuracy data were transformed (McDonald, 2014). Transformed (squared) accuracy values (Table 1) were used for subsequent parametric tests and the terms ‘scene accuracy’, ‘face accuracy’ and ‘size accuracy’ refer to the transformed conditions. Reaction times are reported in Extended Data Table 1-1.

**Table 1.** Oddity task performance. Mean, SD, range and skew for the three conditions accuracy data. Untransformed (left) and squared (right), data are shown. Note that all condition accuracy skews are between −1 and 1 after squaring. See Extended Data Table 1-1 for reaction time data.

| Condition | Mean ± SD (%) | Range | Skew | Squared Mean ± SD (%) | Squared Range | Squared Skew |
| --- | --- | --- | --- | --- | --- | --- |
| Scene | 60.52% ± 7.18 | 38.54-70.83 | -1.05 | 3754.64 ± 807.59 | 1516.90-4973.96 | -0.84 |
| Face | 60.68% ± 8.841 | 39.58-80.21 | -0.08 | 3768.77 ± 1062.13 | 1566.84-6433.38 | 0.40 |
| **Size** | 61.30% ± 13.00 | 29.17-89.58 | -0.16 | 3927.25 ± 1580.89 | 850.70-8025.17 | 0.42 |

### Reduced theta power in the anteromedial hippocampus and posteromedial cortex during scene oddity

First, we sought to examine whether theta power in the hippocampus would be more strongly modulated during scene discrimination trials versus face and size trials (see Fig. 1A, and Materials and Methods).

In the scene>size comparison, there were two negative (i.e., lower theta power in the scene condition compared with the size condition) clusters (cluster p-values = 0.0004; 0.002). These encompassed areas of left medial and inferior temporal lobe (including the hippocampus), and the medial occipital cortex (Figure 1B). Within AAL atlas parcellations, the cluster peaks of theta power reduction were located in the inferior temporal gyrus (MNI coordinates: −46, −16, −36) and cuneus (MNI coordinates: 6, −94, 20). The peak hippocampal theta power reduction was located in the anterior hippocampus (MNI coordinates: −34, −16, −20).

In the face>size comparison, there were two positive (i.e., higher theta power in the face condition compared with the size condition) clusters (cluster p-values = 0.0026; 0.008), encompassing areas of the inferior occipital cortex, and a negative cluster (cluster p-value = 0.0016), encompassing areas of the left lateral and inferior frontal cortices (Figure 1C).

The scene>face comparison revealed a significant negative cluster of theta power reduction (cluster p-value = 0.0002), which encompassed areas of the cerebellum, medial occipital and parietal cortices, including the posteromedial cortex/retrosplenial cortex, and right medial temporal lobe (MTL) areas, including an anteromedial portion of the hippocampus (Figure 1D-E). Within AAL atlas parcellations, the cluster peak of theta power reduction was located in the calcarine cortex (MNI coordinates: −14, −96 −4). The peak hippocampal theta power reduction was located in the anteromedial hippocampus (MNI coordinates: 22 −20 −16; Fig.1E).

### Hippocampal theta power modulation correlates with scene oddity accuracy

We next tested whether hippocampal theta attenuation (see Material and Methods) was related to scene oddity performance. We found modest support for a negative partial correlation (controlling for MEG trial numbers; see Materials and Methods) between hippocampal scene theta and scene oddity accuracy, such that a greater attenuation of theta was associated with greater accuracy (r(37) = −0.331, p = 0.020, 95% CI [−0.508, −0.133], BF10 = 2.41, one-tailed; just exceeding the corrected alpha threshold of 0.017; Figure 2). There was no evidence for significant partial one-tailed correlations between hippocampal theta during the face condition and face oddity accuracy (r(37) = 0.105, p = 0.738, 95% CI [−0.161, 0.296], BF10 = 0.42), or between hippocampal theta during the size condition and size oddity accuracy (r(37) = 0.026, p = 0.563, 95% CI [−0.381, 0.411], BF10 = 0.36). Importantly, the coefficient of the correlation between the scene hippocampal theta power modulation and scene oddity accuracy was significantly stronger than that of the correlation between face hippocampal theta power modulation and face oddity accuracy (z(37) = −2.46, p = 0.007), and that of the correlation between size hippocampal theta power modulation and size oddity accuracy (z(37) = −2.206, p = 0.020).

**Figure 2.**
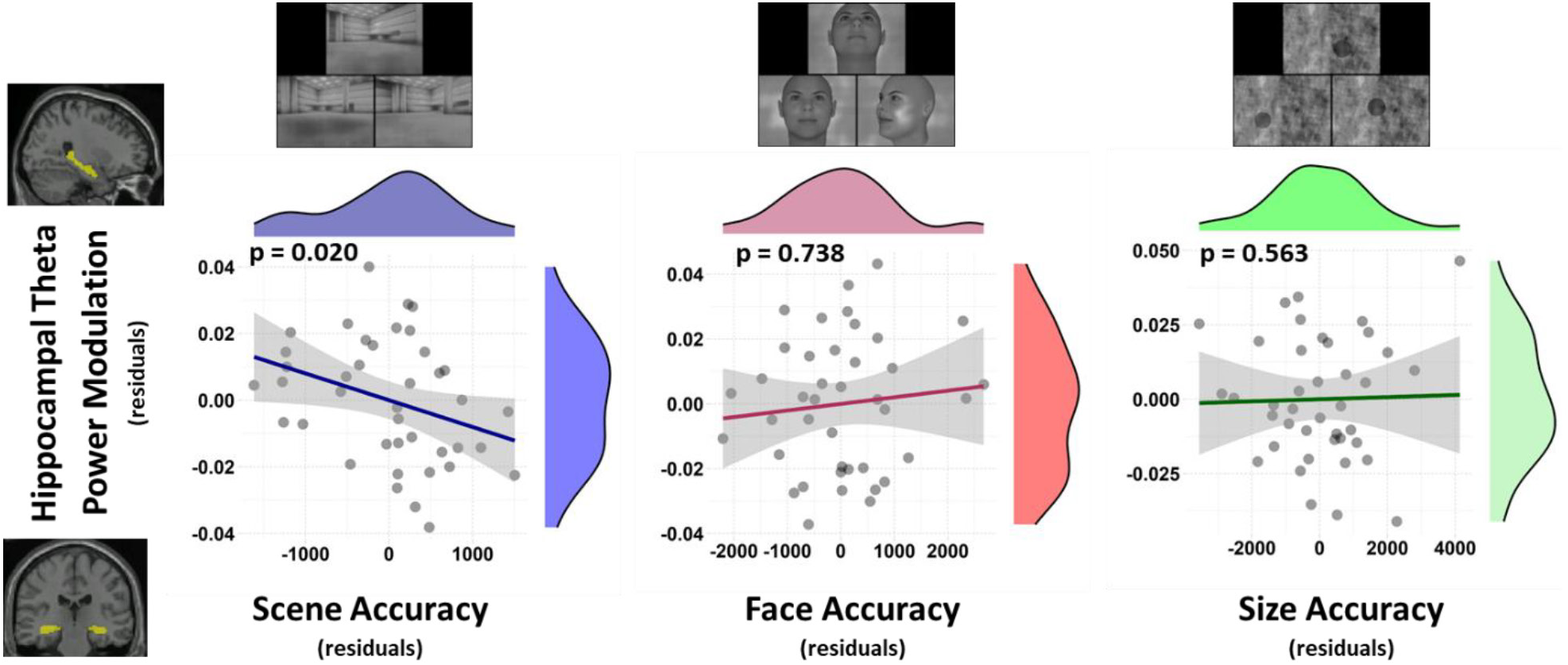
Relationships between oddity task accuracy and hippocampal theta power difference between task and fixation, controlling for the number of MEG trials. The histogram outlines show the distributions of the variables. Note that ‘hippocampus theta power modulation’ refers to modulation during the respective conditions (scene, face and size). The blue/red/green lines are the regression lines, and surrounding shaded areas represent the standard two-tailed 95% confidence interval. Note that these are plots of partial correlations, controlling for the number of MEG trials. Examples of the trials are shown next to the relevant scatter plots. The hippocampus ROI is shown in yellow on a template brain. N = 40.

### Using PCA to identify major features of white-matter tract microstructure

Next, we examined the relationship between tract microstructure (fornix, ILF and PHCB) and oddity performance. To derive major features of white-matter microstructure, we applied advanced biophysical models (i.e., CHARMED and NODDI) as well as the free-water corrected diffusion tensor model (‘FWE-DTI’; see Materials and Methods). These modelling approaches resulted in seven measures per tract (averaged across hemispheres) for each participant (summary statistics shown in Extended Data Figure3-1): FWE-FA; FWE-MD; FWE-Radial Diffusivity (RD); Restricted Fraction (FR); Molecular Protein Fraction (MPF); Neurite Density Index (NDI); and Orientation Dispersion (OD). This larger metric space was then reduced through PCA, a technique that has been shown to be effective in capturing biologically informative features of white-matter microstructure (Chamberland et al., 2019; Geeraert et al., 2020; Henriques et al., 2023; Read et al., 2023). The results from the PCA (KMO: 0.66, sphericity: p<0.0001; Figure 3) showed that 94% of the microstructure data variance was accounted for by the first two principal components, PC1 and PC2. PC1 accounted for 56% of the variance with MD and RD providing the major negative contributions, while FR and MPF provided the major positive contributions (FA: 0.16, MD: −0.48, RD: −0.50, FR: 0.42, MPF: 0.48, OD: 0.16, NDI: −0.23). PC2 accounted for 38% of the variance, with FA and NDI providing the major negative contributions, while OD provided a major positive contribution (FA: −0.56, MD: - 0.17, RD: −0.01, FR: −0.32, MPF: 0.12, OD: 0.55, NDI: −0.48).

**Figure 3.**
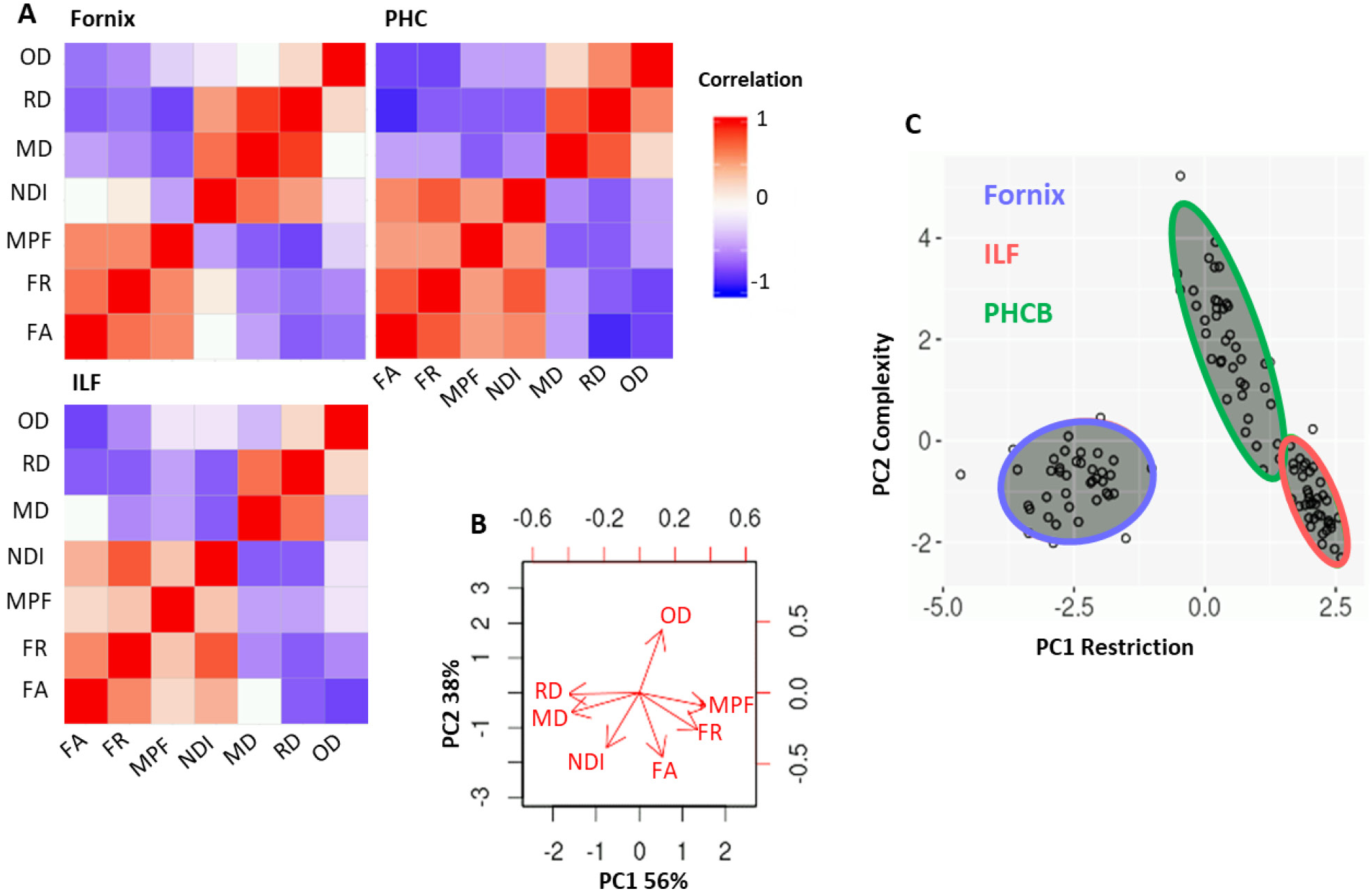
Redundancy between tract microstructure values and results from PCA. **A**) Pearson’s correlations within the microstructure data from each tract suggest that the values give overlapping information. Colour denotes r value according to the key. **B**) Biplot illustrating the influence of each of the measures on PC1 and PC2, which account for 56% and 38% of the variance, respectively. **C**) Tract component scores for each participant, illustrating the differing properties of the tracts. Note that DTI-derived measures are free water corrected (from FWE-DTI). FA, Fractional Anisotropy; FR, Restricted Fraction; ILF, Inferior Longitudinal Fasciculus; MD, Mean Diffusivity; MPF, Macromolecular Protein Fraction; NDI, Neurite Density Index; OD, Orientation Dispersion; PC, Principal Component; PHCB, Parahippocampal Cingulum Bundle; RD, Radial Diffusivity. See Extended Data Table Figure 3-1 for group means and standard deviations for each microstructure value, for each tract.

PC1 was named the ‘tissue restriction’ property of the fibre (the proclivity for water movement along the fibres as opposed to other dispersed directions, likely influenced by axonal packing and myelination) because RD and MPF have been shown to relate to myelin proportion (Kisel et al., 2022; Song et al., 2003; Song et al., 2005), and FR gives an estimate of axon density (De Santis et al., 2014). PC2 was named the ‘microstructural complexity’ property of the fibre (modelled fibre orientation dispersions, likely influenced by underlying tract fanning and crossing), because OD is higher in tracts known to have more fibre fanning and crossing, and it typically correlates more strongly with FA than does NDI (e.g., Zhang et al., 2012), suggesting that fibre orientations influence FA more than the tissue properties themselves. Note that PC1 and PC2 were sign-flipped to aid interpretation, such that increases in PC1 and PC2 reflected increases in tissue restriction and microstructural complexity, respectively.

### The relationship between tissue microstructure and oddity accuracy

As predicted based on Hodgetts et al. (2015), there was a positive correlation between fornix tissue restriction (PC1) and scene oddity accuracy (r(37) = 0.321, p = 0.023, 95% CI [-0.002, 0.564], BF10 = 2.05; Figure 4; just exceeding the corrected alpha threshold of 0.017). The correlations between fornix tissue restriction and face (r(37) = 0.243, p = 0.068, 95% CI [-0.101, 0.537], BF10 = 0.94) or size (r(37) = −0.035, p = 0.585, 95% CI [-0.313, 0.255], BF10 = 0.36) oddity accuracies were not significant. The coefficient of the correlation between fornix tissue restriction and scene oddity accuracy was significantly larger than that of the correlation between fornix tissue restriction and size oddity accuracy (z(36) = 2.052, p = 0.020), but not larger than that of the correlation between fornix tissue restriction and face oddity accuracy (z(36) = 0.446, p = 0.328).

**Figure 4.**
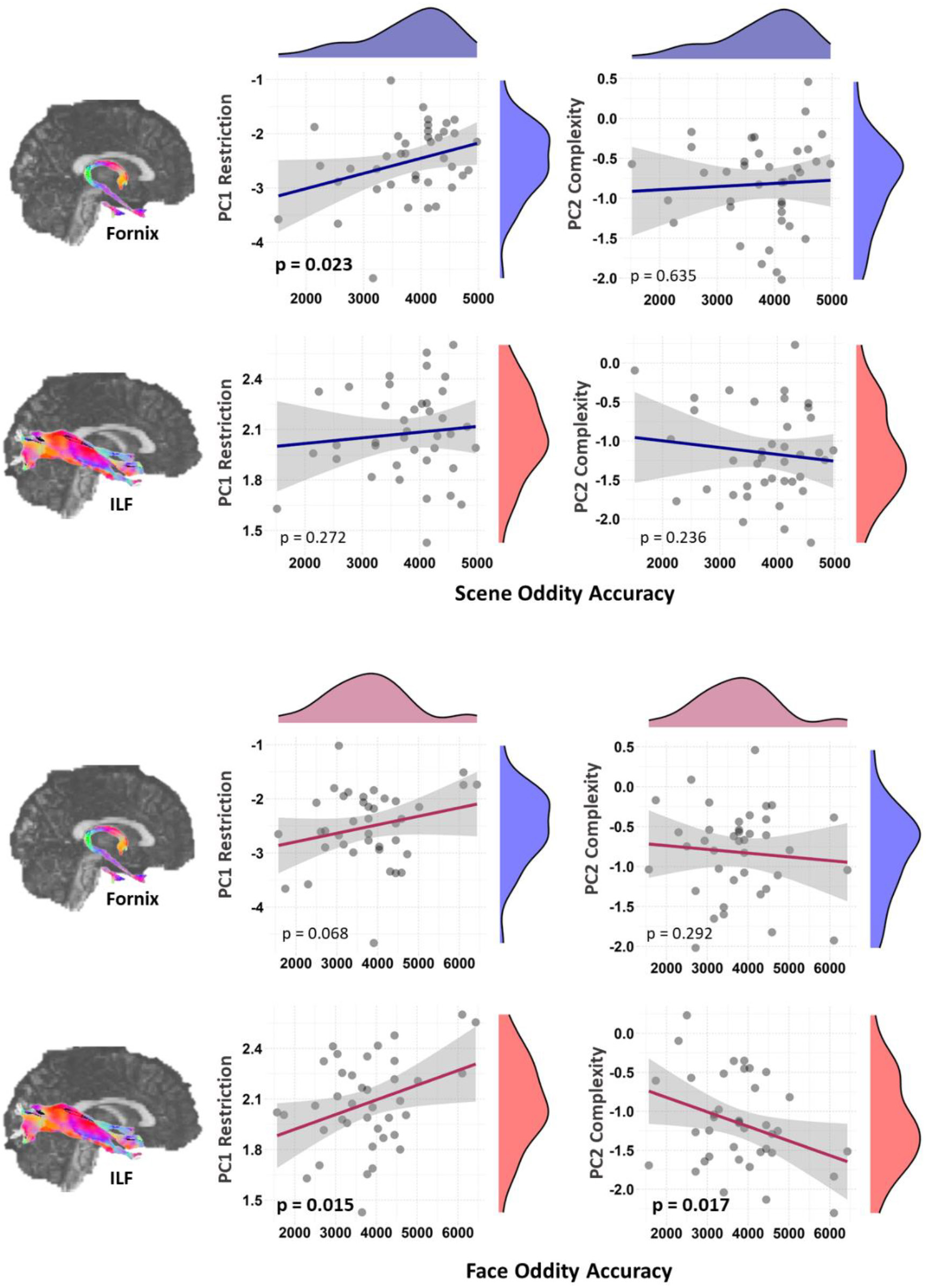
Scatterplots showing the relationship between fornix and ILF microstructure, and scene and face oddity accuracies. The histograms’ outlines show the distributions of the variables. The blue and red lines are the regression lines and surrounding shaded areas represent the standard two-tailed 95% confidence interval. Example images of the tract streamlines (upper: fornix, lower: ILF; standard red-green-blue direction code) are shown next to the relevant axes. N=39. ILF, Inferior Longitudinal Fasciculus; PC, Principal Component. Brain images made from the ExploreDTI example data.

There were no significant correlations between fornix complexity (PC2) and any oddity accuracies (scene (r(37) = 0.057, p = 0.635, 95% CI [-0.222, 0.309], BF10 = 0.38); face (r(37) = −0.090, p = 0.292, 95% CI [-0.419, 0.237], BF10 = 0.41); size (r(37) = 0.088, p = 0.702, 95% CI [-0.236, 0.395] , BF10 = 0.40).

Again as predicted, ILF tissue restriction significantly correlated with face oddity accuracy (r(37) = 0.349, p = 0.015, 95% CI [0.022, 0.590], BF10 = 2.84; Figure 4) and not with scene (r(37) = 0.100, p = 0.272, 95% CI [-0.254, 0.406], BF10 = 0.42) or size (r(37) = −0.037, p = 0.589, 95% CI [- 0.301, 0.229], BF10 = 0.36) oddity accuracy. ILF complexity significantly (inversely) correlated with face oddity accuracy (r(37) = −0.341, p = 0.017, 95% CI [-0.600, −0.002], BF10 = 2.58) and not with scene (r(37) = −0.119, p = 0.236, 95% CI [-0.447, 0.261], BF10 = 0.45) or size (r(37) = 0.170, p = 0.849, 95% CI [-0.106, 0.416], BF10 = 0.57) oddity accuracies. The correlation coefficient between ILF tissue restriction (PC1) and face oddity accuracy was significantly larger than that of the correlation between ILF tissue restriction and size oddity accuracy (z(36) = 2.00, p = 0.023), but was not larger than that of the correlation between ILF tissue restriction and scene oddity accuracy (z(36) = 1.402, p = 0.081). Additionally, the negative correlation coefficient between ILF complexity (PC2) and face oddity accuracy was significantly greater than that of the correlation between ILF tissue restriction and size oddity accuracy (z(36) = −2.722, p = 0.003, but was not greater than that of the correlation between ILF complexity and scene oddity accuracy (z(36) = - 1.250, p = 0.106).

Additionally, we examined the link between PHCB microstructure and oddity performance. In the monkey, retrosplenial fibres join the PHCB to reach parahippocampal areas TH and TF, as well as the presubiculum, parasubiculum, and parts of entorhinal cortex (Bubb et al., 2018). In contrast to the fornix, however, there is little evidence that the PHCB plays a critical role in spatial memory and navigation (Bubb et al., 2018).

Unlike our fornix findings, neither PHCB tissue restriction or complexity correlated with scene oddity accuracy (tissue restriction: r(37) = 0.054, p = 0.372, 95% CI [-0.256, 0.333], BF10 = 0.37; complexity: r(37) = −0.030, p = 0.427, 95% CI [-0.328, 0.307], BF10 = 0.36). Nor did PHCB restriction or complexity correlate with face oddity accuracy or size oddity accuracy (face oddity accuracy - tissue restriction: r(37) = 0.294, p = 0.034, 95% CI [-0.098, 0.583], BF10 = 1.52; complexity: r(37) = −0.258, p = 0.057, 95% CI [-0.528, 0.051], BF10 = 1.07). size oddity accuracy - tissue restriction: r(37) = 0.180, p = 0.136, 95% CI [-0.158, 0.474], BF10 = 0.61; complexity: r(37) = −0.106, p = 0.260, 95% CI [-0.387, 0.216], BF10 = 0.43).

Multiple linear regression was used to assess whether fornix microstructure predicted scene oddity accuracy beyond ILF and PHCB microstructure. A model with fornix tissue restriction, PHCB tissue restriction and ILF tissue restriction did not significantly predict scene oddity accuracy (adjusted R^2^ = 0.029, p = 0.267) but fornix tissue restriction (PC1) was an independent predictor (p = 0.031, adjusted for one-tailed prediction). These results indicate that microstructure of the fornix, specifically the ‘tissue restriction’ component, relates to scene processing performance, whereas microstructure properties of the ILF and PHCB do not.

### No evidence for an association between fornix tissue restriction and hippocampal theta during scene accuracy

A partial correlation test (controlling for number of trials) showed that fornix tissue restriction and scene hippocampal theta power modulation did not significantly correlate (r(36) = - 0.078, p = 0.642, 95% CI [-0.360, 0.270], BF10 = 0.39), despite both having been found to relate to scene oddity accuracy. Multiple linear regression was used to assess whether fornix tissue restriction and scene hippocampal theta power modulation influences on scene processing were independent of one another. In a model with scene oddity accuracy as the dependant variable, and fornix tissue restriction, scene hippocampal theta power modulation and scene MEG trials numbers^1^ as covariates (adjusted R^2^ = 0.225, p = 0.007), fornix tissue restriction and scene hippocampal theta power modulation were both independent predictors (p-values = 0.035, 0.034, standardized regression coefficients = 0.269, −0.270, respectively).

### No evidence that the findings purely reflect incidental encoding

Lastly, to test if our findings of relationships between scene discrimination performance and scene hippocampal theta modulation / fornix tract microstructure were influenced by incidental encoding, we examined relationships between these imaging measures and performance in the subsequent surprise memory task.

Overall, memory performance was poor, which was understandable as the participants were not told to memorize the oddity stimuli. The mean d’ scores for scenes and faces were 0.49 (SD = 0.452) and 0.28 (SD = 0.287), respectively. There was no significant partial correlation (controlling for the number of oddity scene MEG trials) between scene oddity hippocampal theta power modulation and scene d’ (r(37) = 0.171, p = 0.298, 95% CI [-0.186, 0.515], BF10 = 0.57).

Also, there was no significant correlation between scene d’ and fornix PC1 (r(37) = 0.137, p = 0.203, 95% CI [-0.223 0.483], BF10 = 0.48; one-tailed).

## Discussion

Here, we advanced understanding of the role of an extended hippocampal (Aggleton, 2012) or posteromedial (Murray et al., 2017; Ranganath & Ritchey, 2012) system in cognition by demonstrating theta power attenuation, during scene compared with face or shape- size discrimination, in the hippocampus alongside the parahippocampal and posteromedial cortices. Furthermore, inter-individual differences in scene hippocampal theta power attenuation correlated with scene, relative to face, oddity accuracy. There was also a correlation between scene oddity accuracy and fornix microstructure, specifically the ‘tissue restriction’ property.

Conversely, ILF microstructure correlated with face, but not scene oddity accuracy.

### Scene discrimination may be supported by theta power modulation in posteromedial cortex, including an anteromedial hippocampus scene-selective ‘hub’

When intersecting our scene-related theta modulation maps, we identified a peak in hippocampal theta modulation in the comparison of scene versus face oddity in the anteromedial hippocampus, aligning with previous fMRI studies of scene imagery (Zeidman et al., 2015) and scene discrimination (Hodgetts, Voets, et al., 2017). High-resolution fMRI work suggests this scene-selective region likely corresponds to the anteromedial subicular complex (Hodgetts, Voets, et al., 2017; Read et al., 2024), which connects the hippocampus, (primarily via the fornix) to the extended hippocampal system (Aggleton, 2012), and which receives spatial input from cortical areas including retrosplenial and parahippocampal cortices (Aggleton, 2012; Kravitz et al., 2011). Therefore, our results may reflect the scene-selective processes taking place in the subicular complex, specifically. Relatedly, previous source-space MEG studies revealed reduced anterior hippocampal theta power during scene imagery (Barry et al., 2019). The clusters of reduced theta power during scene oddity also included extra-hippocampal MTL, medial occipital, and posteromedial cortices. Prior research has identified reduced theta power across these regions related to scene imagery (Barry et al., 2019) and spatial memory (Crespo-Garcia et al., 2016; Fellner et al., 2016). Collectively, these results suggest that reduced theta power in subicular complex and interconnected cortical regions may reflect a commonality of processes across these tasks, such as mental scene construction (Zeidman & Maguire, 2015).

Although the neuronal bases of reduced theta power are unclear, our finding of a relationship between theta power reduction and scene oddity accuracy indicates that such reduced theta power reflects neuronal processes beneficial to scene discrimination. Hanslmayr et al. (2012) argued low-frequency power decreases are a mechanism to de-correlate neural activity to enhance regional neural coding capacity. Although originally formulated in the context of alpha and beta power reduction during memory encoding, the principles may also extend to hippocampal theta.

### Fornix tissue restriction linked to scene discrimination

Considering that most subiculum outputs rely on the fornix (Aggleton, 2012), it is unsurprising that the fornix has been implicated in scene-based cognition (Gaffan, 1994). Hodgetts et al. (2015) found fornix MD and ILF MD to correlate with scene oddity and face oddity accuracy, respectively. Here, we link scene discrimination to more specific features of fornix microstructure. Redundancies in several microstructure measures were exploited to reveal two biologically interpretable components, comparable to those in previous studies (Chamberland et al., 2019; Read et al., 2023). PC1 was most influenced negatively by MD, RD and positively by FR and MPF, and was interpreted as positively relating to a ‘tissue restriction’ (axonal packing/myelination) property of the fibre. PC2 was most influenced positively by OD and negatively by FA and was interpreted as negatively relating to a ‘microstructural complexity’ (tract fanning/crossing) property of the fibre (Chamberland et al., 2019). We found a relationship between fornix tissue restriction and scene discrimination, whereas both ILF microstructure components (tissue restriction and complexity) correlated with face oddity discrimination. Linear regression analysis indicated fornix tissue restriction as an independent microstructural predictor (from ILF and PHCB tissue restriction) of scene discrimination. These findings align with nonhuman primate work showing spatial scene memory deficits following fornix, but not cingulum lesions (Gaffan, 1994; Parker & Gaffan, 1997).

### Fornix fibre restriction and hippocampal theta modulation independently influence scene processing

The fornix is a key conduit for theta in the extended hippocampal system, conveying hippocampal inputs from the medial septum (involved with theta oscillation generation; Rawlins et al., 1979), as well as hippocampal outputs to the anterior thalamus (containing theta modulated cells; Jankowski et al., 2013). However, we found no evidence that the relationship between fornix microstructure and scene discrimination was mediated by hippocampal theta power modulation. Rather, fornix tissue restriction and scene hippocampal theta power modulation were independent predictors of scene discrimination.

This echoes previous findings (Hodgetts et al., 2015) that hippocampal BOLD did not mediate the relationship between fornix MD and scene discrimination. Together, these findings suggest that fornix microstructure and hippocampal activity independently influence scene discrimination. In rodents, while fornix and hippocampal lesions frequently result in comparable spatial deficits (Aggleton & O’Mara, 2022; Aggleton et al., 2010), dissociations between fornix and hippocampus lesions have been reported (Dumont et al., 2015). Non-fornical cortical connectivity, presumably important to hippocampal scene activity, includes connections between subiculum and retrosplenial cortex (Aggleton, 2012). Retrosplenial cortex, alongside the hippocampus, was included in the posteromedial cluster of theta power reduction during scene discrimination and is known to support multiple aspects of spatial processing (Vann et al., 2009). Our findings of independent contributions of fornix microstructure and hippocampal/cortical theta align with recent proposals of partially independent hippocampal-cortical and medial diencephalic-cortical processing streams (Aggleton & O’Mara, 2022).

### Face discrimination is supported by ILF microstructure

The ILF interconnects occipital and anterior temporal cortices including perirhinal cortex (Herbet et al., 2018), a region critical to face oddity discrimination (Barense et al., 2007). ILF properties have been related to face discrimination (Hodgetts et al., 2015), facial recognition (Behrmann et al., 2007) and face-naming (Burkhardt et al., 2023). Unlike the fornix, where tissue restriction appeared more important to scene discrimination, ILF tissue restriction and complexity both correlated with face oddity accuracy, positively and negatively, respectively, the latter of which may reflect the branching patterns or crossing of axons.

## Limitations

Our study has some limitations. First, there is a necessary discrepancy in how trials with no response were treated in the MEG (excluded) and behavioural data (classed as incorrect) which was addressed by using partial correlations (controlling for MEG trial numbers). The inclusion of unanswered trials in the MEG data processing may have reduced sensitivity to task relevant signals as it is not clear if the participant was distracted, but in the behavioural data, if unanswered trials are removed then, when calculating the percentage of correct responses, scores of participants who missed trials would be inflated, as we can assume that either prolonged trial attempts or distraction would likely result in an incorrect answer. Importantly, there were no correlations between MEG- measured task oscillatory power differences and MEG trial numbers.

Second, we interpret our MEG results mostly considering previous fMRI work and it should be noted that spatial location in MEG is less accurate than that of fMRI. Although it has previously been assumed that the hippocampus is too deep a structure to measure with MEG (see Pu et al., 2018 for review), it is understood that the distance to the sensors is partly counteracted by the high source-current density generated by the pyramidal cell layers of the hippocampus which is more than twice that of neocortical grey matter (Attal & Schwartz, 2013; Ruzich et al., 2019), and hippocampal signal detection with MEG has been validated using concurrent intracranial electrophysiology (iEEG) and modelling (e.g., Attal & Schwartz, 2013; Pizzo et al., 2019). Moreover, despite deep source localization challenges, we are confident our results demonstrate hippocampal contribution, rather than just signal spread from nearby MTL cortex. First, multiple studies have successfully measured hippocampal signals with MEG (Barry et al., 2019; Meyer et al., 2017; Mills et al., 2012; Pu et al., 2018) whereas few report perirhinal cortex signals (e.g., Moses et al., 2009). Second, we found that hippocampal theta power modulation during scene oddity correlated with discrimination ability, consistent with the effects of hippocampal lesions that spare parahippocampal cortex (Lee, Bussey, et al., 2005). Also relating to spatial location accuracy of MEG, the apparent unilateral hippocampal affects in our results should be interpreted with caution. Beamforming techniques assume sources are uncorrelated, supressing bilateral responses, which may explain cases where fMRI and MEG studies find bilateral and unilateral effects, respectively, despite using the same kind of behavioural tasks (O’Neill et al., 2021).

Regarding the microstructure analyses, it should be noted that, while a PCA technique was applied to increase biological interpretability of microstructure features, tractography does not capture fibre properties directly, and interpretations of PCA components have not been confirmed histologically (see Read et al., 2023 for discussion).

## Conclusions

In summary, we found specific engagement of an extended hippocampal system during scene discrimination, in the form of modulated theta power activity, and that hippocampal theta power modulation correlated with scene oddity performance. Moreover, fornix tissue restriction was important for scene oddity performance, independent of hippocampal theta. Conversely, ILF microstructure correlated with face oddity performance. Together, this work provides novel support for multiple-system, representation-based accounts of the MTL (Aggleton & O’Mara, 2022; Murray et al., 2017; Ranganath & Ritchey, 2012).

## Additional information

## Acknowledgements

We would like to thank Dr Gavin Perry (Cardiff University) for his help with MEG protocol set-up and advice. This work was supported by the Biotechnology and Biological Sciences Research Council (BBSRC) [BB/V010549/1; BB/V008242/2; to C.J.H., A.D.L., K.S.G], a Wellcome

Institutional Strategic Support Fund award to A.D.L, and Cardiff University School of Psychology PhD studentship to M-L.R.

## Data sharing

The raw neuroimaging data cannot be shared publicly due to GDPR (General Data Protection Regulation) -related ethical restrictions (pertaining to the study’s participant consent form) which do not allow for the public archiving of study data. However, access to pseudo-anonymized data could be granted following signing and approval of suitable data-transfer agreements. Readers seeking access should contact the corresponding author. Analysis code will be made available.

## CRediT author statement

Conceptualization: K.U.-R., M-L.R, K.S.G., and K.D.S. Data curation: M-L.R. Formal analysis: M-

L.R. Funding acquisition: K.S.G., K.U.-R., K.D.S., A.D.L. Investigation: M-L.R. Project administration: K.U.-R., M-L.R. Software: C.J.E. Supervision: K.U.-R., K.S.G., K.D.S., A.D.L. and

C.J.H. Visualization: M-L.R. Writing - original draft: M-L.R., K.U.-R. K.S.G., K.D.S. and C.J.H.

Writing - review & editing: M-L.R., K.U.-R. K.S.G., K.D.S., C.J.E, A.D.L. and C.J.H.

## Extended Data

**Figure 3-1.**
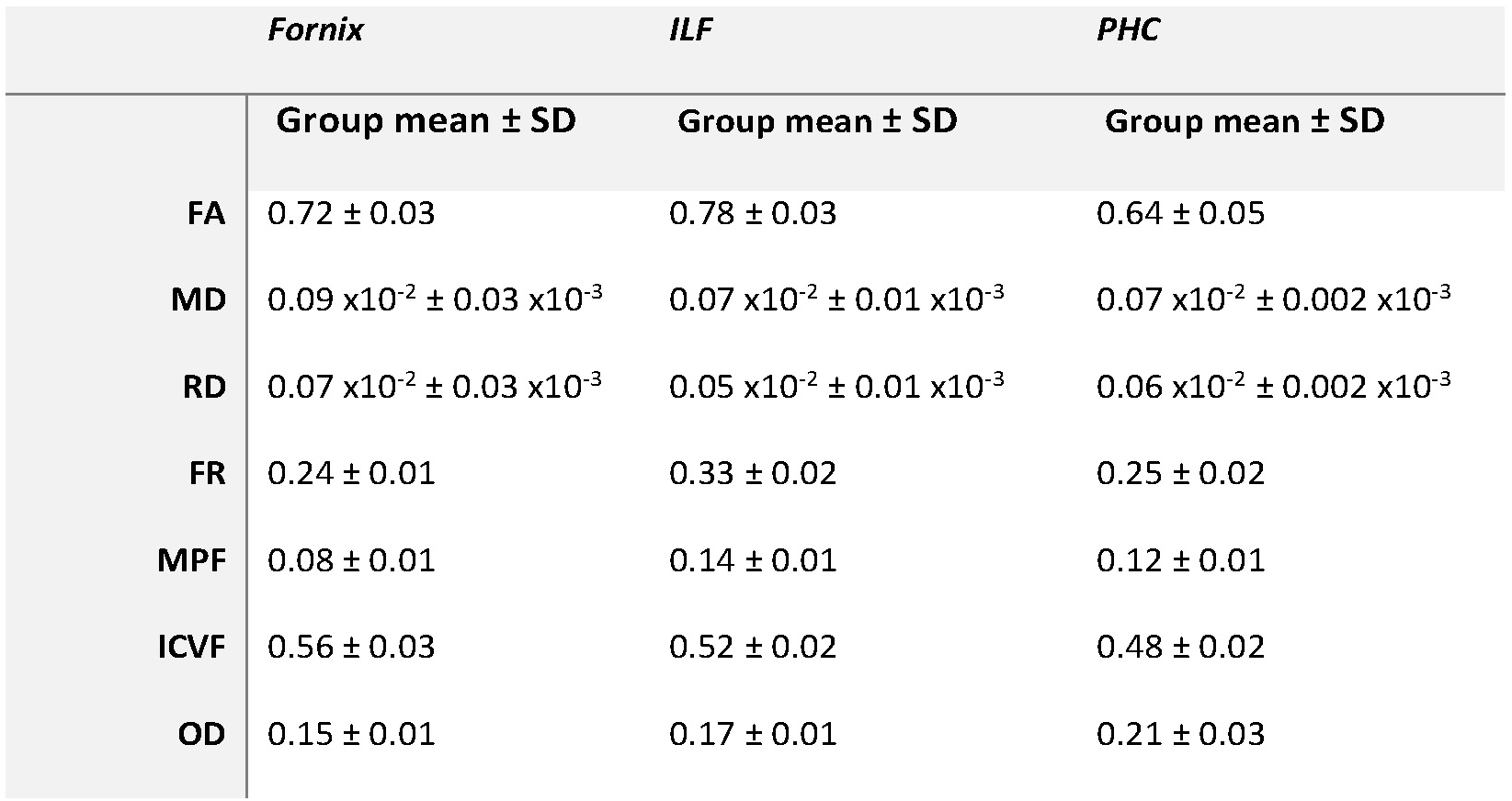
Group mean and standard deviation for each microstructure value, for each tract. Microstructure values were averaged over tract streamlines for each participant. FA: Fractional Anisotropy. FR: Restricted Fraction. ICVF: Intracellular Volume Fraction. MD: Mean Diffusivity. MPF: Macromolecular Proton Fraction. OD: Orientation Dispersion. RD: Radial Diffusivity.

**Table 1-1.** Oddity task reaction times. Mean, SD, range and skew for the three conditions reaction time data.

| Condition | Mean ± SD (s) | Range | Skew |
| --- | --- | --- | --- |
| Scene | 5.14 ± 0.51 | 3.95 - 6.00 | -0.81 |
| Face | 4.16 ± 0.78 | 2.56 - 5.48 | -0.06 |
| Size | 3.14 ± 0.70 | 2.14 - 4.96 | 0.78 |

## Footnotes

1 Included to control for the number of trials, see Materials and Methods.

